# Reply to: Structural polymorphism of amyloid fibrils in ATTR amyloidosis revealed by cryo-electron microscopy

**DOI:** 10.64898/2026.10.01.755719

**Authors:** Candace Kelly, Binh An Nguyen, Virender Singh, Preeti Singh, Parker T. Bassett, Saylor Schreiber, Christian Lopez Escobar, Barbara Kluve-Beckerman, Bret M. Evers, Lorena Saelices

## Abstract

In the paper by Nguyen et al, published in *Nature Communications* in 2024^1^, we describe structural polymorphism in fibrils extracted from patients with ATTR amyloidosis. ATTR amyloidosis is characterized by deposition of transthyretin amyloid fibrils across multiple organs, resulting in marked phenotypic variability. Previous studies on ATTR suggest its phenotypic variability may be linked to differences in fibril structure, as demonstrated in amyloid diseases such as AA amyloidosis, tauopathies, or synucleinopathies^2–4^. Thus far, most ATTR fibrils adopt a strikingly similar fold across patients and affected organs^5–8^.

The most notable exception is described in the study by Nguyen et al., where we report the identification of two co-existing polymorphs in cardiac fibrils from three patients with the ATTRv-I84S variant. These cardiac polymorphs differ in conformation of the segment Gly57 to Gly67, termed the *gate*, which regulates access to a central polar channel within the fibril core. In one polymorph, the gate adopts an open conformation, exposing the channel, whereas in the other it is closed, shielding it^1^. Whether these polymorphs extend beyond cardiac tissue remains unknown. Here, we address this question by extending our analysis to additional organs from a fourth patient with ATTRv-I84S amyloidosis.

## ATTRv-I84S fibril polymorphism extends across organs

We studied the heart, spleen, and kidney from an additional ATTRv-I84S patient. We performed tissue staining, fibril extraction, mass spectrometry, and structure determination, as permitted by tissue availability, enabling multi-level characterization (Figure 1, and Supplementary Table 1). Histological staining confirmed amyloid deposition in the spleen and kidney (Figure 1a and Supplementary Figure 1a); insufficient heart tissue was available for histological analysis because the sample was prioritized for cryo-EM structure determination. We extracted amyloid fibrils from all tissues using a mild, water-based method to minimize structural perturbation, and assessed fibril morphology and sample quality by transmission electron microscopy (Figure 1b). We confirmed the presence of both wild-type and variant transthyretin in the heart, spleen, and kidney using mass spectrometry (Figure 1c), consistent with the heterozygous genotype.

**Figure 1:**
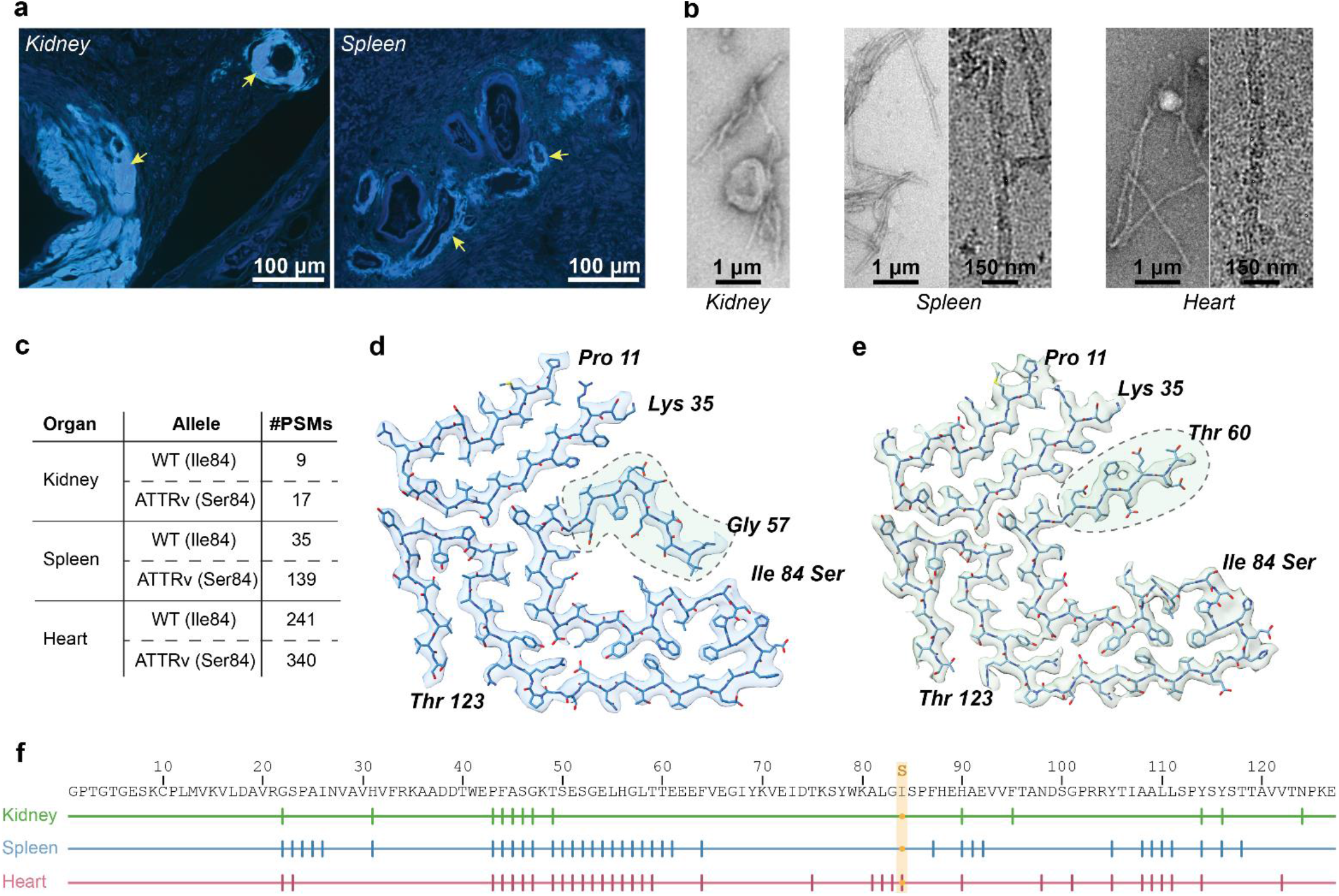
Characterization of amyloid fibrils from kidney, spleen and heart in an ATTRv-I84S amyloidosis patient. **a**, Thioflavin S staining of kidney and spleen tissue sections confirms the presence of amyloid deposits (arrows). **b**, Representative transmission electron microscopy images of fibrils extracted from kidney, spleen, and heart tissue. Representative cryo-EM micrographs from spleen and heart fibrils are shown on the right. **c**, Tryptic mass spectrometry analysis of fibrils extracted from heart, spleen, and kidney confirms that the patient is heterozygous for the ATTRv-I84S mutation. Protein-spectrum matches with high confidence outputs are shown. **d, e**, Representative cryo-EM density maps and atomic models of fibrils extracted from the **s**pleen, revealing closed-gate (**d**) and open-gate (**e**) folds previously described in Nguyen *et al*. (*Nature Communications*, 2024)^1^. Dashed circles highlight the gate region. **f**, Proteolytic profiles of fibrils extracted from the indicated organs. Residues identified by mass spectrometry are shown below the transthyretin sequence. Colors indicate the corresponding tissue source.

In agreement with our prior observations, we identified two distinct fibril folds in heart and spleen tissues by cryo-EM, corresponding to the previously described open- and closed-gate morphologies. The structural differences were primarily localized to the gate encompassing residues Gly57 to Gly67. In the closed-gate fold, this segment occludes the polar channel, similar to that observed in wild-type ATTR fibrils (Figure 1d and Supplementary Figure 1b), whereas in the open-gate fold it reorients outward, exposing a solvent-accessible pocket (Figure 1e and Supplementary Figure 1c). Both polymorphs can co-exist within the same fibril (Supplementary Figure 1e). In contrast, the yield of fibril extracted from the kidney was insufficient for cryo-EM reconstruction. These results show that polymorphism in ATTRv-I84S is not restricted to the heart but extends to the spleen, suggesting that polymorphism may also occur in other organs.

We observed comparable proteolytic cleavage patterns across three organs, consistent with those reported in other ATTR variants (Figure 1f)^5–8^. These patterns suggest that proteolysis may occur both before and after fibril formation. Several cleavage sites within the ordered core are not accessible in the fibrillar state, supporting cleavage prior to assembly, whereas multiple sites within the disordered Ala36 to His56 region remain accessible in the fibrillar state, indicating that cleavage can also occur after fibril formation.

The persistence of the same fibril folds in heart and spleen tissue suggests that this structural conservation is established before fibrils are distributed across organs. This could occur through aggregation in the circulation before organ deposition or through initial fibril formation in one organ followed by seeding-mediated spread to other tissues^9^. Although the mutation may govern formation of specific polymorphs, diverse biochemical environments of different tissues would be expected to produce structural divergence if fibrils formed independently. Instead, these mutation-driven folds persist across tissue environments. In agreement with this model, previous detection of circulating transthyretin aggregates in patients with both wild-type and variant ATTR amyloidosis supports systemic initiation of aggregation prior to deposition^9^. Defining the mechanisms that initiate aggregation in the circulation will be critical to understanding early disease pathogenesis.

This study has limitations. Most patients analyzed belong to the same kindred, and validation in independent cohorts is necessary. Limited tissue availability precluded histochemistry of the cardiac extract, while the limited yield of fibrils extracted from the kidney prevented cryo-EM structural determination.

## Acknowledgements

We thank the Structural Biology Laboratory, the UTSW Cryo-Electron Microscopy Facility, the UTSW Electron Microscopy Core Facility (partially supported by CPRIT grant RP220582), the UTSW Proteomics Core, and the UTSW Histo Pathology Core (supported in part by the Leducq Foundation for Cardiovascular Research, CPRIT, and the NIH), as well as the national cryo-EM facilities at Stanford-SLAC (supported by NIGMS 1R24GM154186). Computational resources were provided by the BioHPC supercomputing facility (https://portal.biohpc.swmed.edu) and UCSF ChimeraX (Resource for Biocomputing, Visualization, and Informatics, UCSF; supported by NIH R01-GM129325 and the NIAID Office of Cyber Infrastructure and Computational Biology). The content is solely the responsibility of the authors and does not necessarily represent the official views of the NIH.

ChatGPT (OpenAI, 2026) was used to help edit and improve the clarity of the manuscript; the authors reviewed and verified all AI-assisted content.

This work was supported by the American Heart Association (Career Development Award 847236, L.S.); the National Institutes of Health, National Heart, Lung, and Blood Institute (New Innovator Award DP2-HL163810 and R01-HL177670, L.S.); the Welch Foundation (Research Award I-2121-20220331, L.S.); and a UTSW Endowment (Distinguished Researcher Award from the President’s Research Council and start-up funds, L.S.).

## Author contributions

Conceptualization: L.S., B.A.N., C.K. Methodology: L.S., B.A.N., V.S. Investigation: B.A.N., V.S., P.S., C.K., P.B., C.L.E., B.K.B., B.E., L.S. Visualization: B.A.N., C.K., P.B., S.S., L.S. Supervision: B.A.N, L.S. Funding acquisition: L.S. Project administration L.S. Writing – original draft: C.K., B.A.N., L.S., writing – review & editing: C.K., B.A.N., L.S.

## Competing interests

L.S. and B.A.N. are co-founders of AmyGo. L.S. reports consulting and/or advisory board fees from Pfizer, AstraZeneca, and AmyGo, and research support from the NIH, AstraZeneca, and UT Southwestern Medical Center. These relationships had no influence on the design, conduct, or reporting of the present study. The remaining authors declare no competing interests.

## Notes

ftp://

## References

1. Nguyen, B. A. et al. Structural polymorphism of amyloid fibrils in ATTR amyloidosis revealed by cryo-electron microscopy. Nat. Commun. 15, 581 (2024).

2. Banerjee, S. et al. Amyloid fibril structure from the vascular variant of systemic AA amyloidosis. Nat. Commun. 13, 7261 (2022).

3. Liberta, F. et al. Cryo-EM fibril structures from systemic AA amyloidosis reveal the species complementarity of pathological amyloids. Nat. Commun. 10, 1104 (2019).

4. Scheres, S. H. W., Ryskeldi-Falcon, B. & Goedert, M. Molecular pathology of neurodegenerative diseases by cryo-EM of amyloids. Nature 621, 701–710 (2023).

5. Fernandez-Ramirez, M. D. C. et al. A shared amyloid fold in cardiac fibrils from three neuropathy-associated ATTR variants. Structure 34, 487-495.e3 (2026).

6. Nguyen, B. A. et al. ATTRv-V30M amyloid fibrils from heart and nerves exhibit structural homogeneity. Structure 32, 2244-2250.e3 (2024).

7. Nguyen, B. A. et al. Cryo-EM confirms a common fibril fold in the heart of four patients with ATTRwt amyloidosis. Commun. Biol. 7, 905 (2024).

8. Fernandez-Ramirez, M. D. C. et al. Structural and molecular homogeneity of ATTRv-T60A amyloid fibrils across patients and organs. Structure 33, 2013-2019.e4 (2025).

9. Pedretti, R. et al. Detection of Circulating Transthyretin Amyloid Aggregates in Plasma: A Novel Biomarker for Transthyretin Amyloidosis. Circulation 149, 1696–1699 (2024).

