## Supplementary Information for "Reply to: Structural polymorphism of amyloid fibrils in ATTR amyloidosis revealed by cryo-electron microscopy"

**Supplementary Table 1: Experiments conducted on each organ based on availability and/or suitability.**

| | **Heart** | **Spleen** | **Kidney** |
|---|---|---|---|
| **Histology** | No | Yes | Yes |
| **Tryptic Mass Spectrometry** | Yes | Yes | Yes |
| **Extraction** | Yes | Yes | Yes |
| **TEM** | Yes | Yes | Yes |
| **CryoEM & data processing** | Yes | Yes | No |

**Supplementary Table 2: Cryo-EM data collection, refinement and validation statistics.**

| | #1 Heart | | #2 Spleen | |
|---|---|---|---|---|
| | Close (EMDB-78674) (PDB 38AE) | Open (EMDB-78675) (PDB 38AF) | Close (EMDB-78679) (PDB 38AJ) | Open (EMDB-78680) (PDB 38AK) |
| **Data collection and processing** | | | | |
| Magnification | 130,000 | | 130,000 | |
| Voltage (kV) | 300 | | 300 | |
| Electron exposure (e–/Å$^2$) | 40 | | 40 | |
| Exposure time (s) | 4.16 | | 4.16 | |
| Defocus range (μm) | -0.8 to -2.6 | | -0.8 to -2.6 | |
| Pixel size (Å) | 0.954 | | 0.954 | |
| Symmetry imposed | C1 | | C1 | |
| Rise | -1.20 | -1.27 | -1.22 | -1.25 |
| Twist | 4.95 | 4.95 | 4.95 | 4.95 |
| Initial particle images (no.) | 1,323,905 | 1,323,905 | 867,558 | 867,558 |
| Final particle images (no.) | 17,481 | 43,899 | 37,513 | 46,811 |
| Map resolution (Å) | | | | |
| FSC threshold 0.143 | 3.4 | 3.4 | 3.0 | 3.2 |
| Map resolution range (Å) | 3.1 – 4.5 | 3.2 – 4.6 | 2.9 – 3.8 | 2.8 – 4.1 |
| **Refinement** | | | | |
| Initial model used (PDB code) | 8TDN | 8TDO | 8TDN | 8TDO |
| Model resolution (Å) | | | | |
| FSC threshold | 3.6 | 3.4 | 2.6 | 3.0 |
| Map sharpening *B* factor (Å$^2$) | -95.74 | -114.36 | -99.27 | -110.405 |
| Model composition | | | | |
| Non-hydrogen atoms | 3580 (5 chains) | 2788 (4 chains) | 3580 (5 chains) | 2724 (4 chains) |
| Protein residues | 460 | 356 | 460 | 348 |
| Ligands | 0 | 0 | 0 | 0 |
| R.m.s. deviations | | | | |
| Bond lengths (Å) | 0.004 | 0.004 | 0.004 | 0.003 |
| Bond angles (°) | 0.859 | 0.744 | 0.675 | 0.565 |
| Validation | | | | |
| MolProbity score | 1.56 | 1.83 | 1.68 | 1.22 |
| Clashscore | 9.52 | 12.07 | 5.46 | 4.42 |
| Poor rotamers (%) | 0.00 | 0.00 | 0.00 | 0.00 |
| Ramachandran plot | | | | |
| Favored (%) | 97.73 | 96.47 | 94.32 | 98.80 |
| Allowed (%) | 2.27 | 3.53 | 5.68 | 1.20 |
| Disallowed (%) | 0.00 | 0.00 | 0.00 | 0.00 |

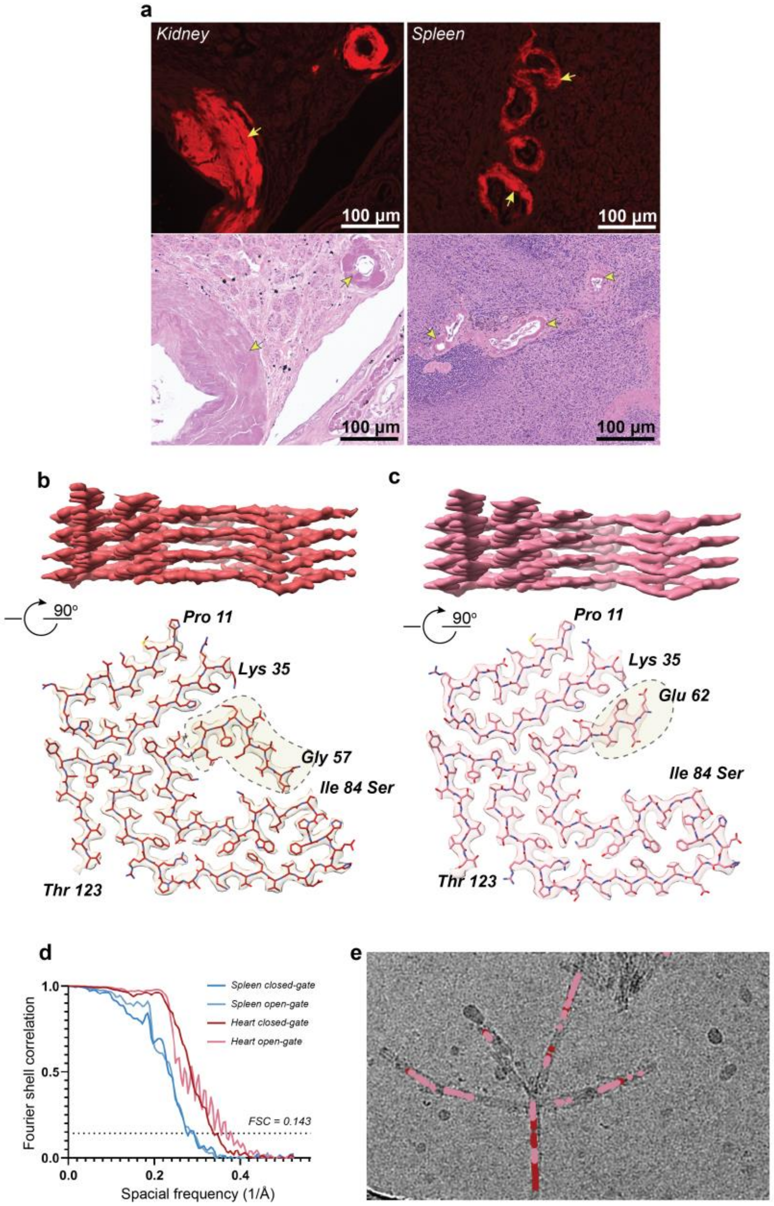


**Supplementary Figure 1: Graphical summary of data. a** Histochemistry using Congo Red (top) and hematoxylin and eosin (bottom) show amyloid deposits in the kidney and spleen. **b** Closed and **c** open gate morphologies are present in heart and spleen 3D classes as seen in Nguyen et al, published in *Nat Comm* 2024[1]. **d** FSC shows the normalized cross-correlation between 3D reconstructions. **e** Tracing maps show that both morphologies can coexist on the same fibril.

### Supplementary Methods

#### Ethical statement

This patient sample was fully anonymized prior to transfer and was therefore exempt from Institutional Review Board oversight by UT Southwestern Medical Center's Office of the Human Research Protection Program. Informed consent for tissue collection and research use was obtained under the protocols governing the original studies conducted at Indiana University School of Medicine. No participants were recruited for the present study, and no compensation was provided.

#### Patient samples and extraction of amyloid fibrils from tissue

We obtained kidney, spleen, and heart tissue postmortem from an ATTR patient carrying I84S mutation from the laboratory of Dr. Merrill D. Benson at the Indiana University School of Medicine. The patient was a female in her mid 50s.

*Ex-vivo* amyloid fibril preparations were derived from fresh-frozen human tissue following previously established protocols[1–5]. Approximately 200 mg of frozen tissue was allowed to thaw at room temperature before being diced into small fragments using a scalpel. The resulting tissue pieces were then resuspended in 1 mL of Tris-calcium buffer (20 mM Tris, 150 mM NaCl, 2 mM $CaCl_2$, 0.1% $NaN_3$, pH 8.0) and pelleted by centrifugation at 3,100×g for 5 minutes at 4 °C. The pellet underwent four additional washes in the same buffer. Following this washing series, the pellet was resuspended in 1 mL of collagenase solution (5 mg/mL, prepared in Tris-calcium buffer) and incubated overnight at 37 °C with continuous shaking at 400 rpm. The suspension was then centrifuged at 3,100×g for 30 minutes at 4 °C, and the resulting pellet was taken up in 1 mL of Tris–EDTA buffer (20 mM Tris, 140 mM NaCl, 10 mM EDTA, 0.1% $NaN_3$, pH 8.0). After a 5-minute centrifugation at 3,100×g and 4 °C, the Tris–EDTA wash was repeated nine more times. All supernatants were retained for downstream analysis where applicable. The washed pellet was subsequently resuspended in 200 µL of ice-cold water containing 5–10 mM EDTA and centrifuged for 5 minutes at 3,100×g and 4 °C; amyloid fibrils released into the supernatant were collected at this stage, with EDTA facilitating fibril solubilization. This extraction cycle was performed 3–5 times in total. All patient-derived material was processed and analyzed independently.

#### Negative-stain transmission electron microscopy

Successful fibril extraction was verified by negative-stain transmission electron microscopy using an established protocol[1–5]. Briefly, 3 µL of sample was deposited onto a glow-discharged carbon film 300-mesh copper grid (Electron Microscopy Sciences) and allowed to adsorb for 2 minutes, after which excess solution was carefully wicked away with filter paper. The grid was then stained by applying 5 µL of 2% uranyl acetate for 2 minutes, followed by blotting to remove the stain. A second 5 µL aliquot of uranyl acetate was immediately applied and removed. Grids were examined using an FEI Tecnai 12 electron microscope operating at an accelerating voltage of 120 kV.

#### Cryo-EM sample preparation, screening, data collection, and processing

Freshly isolated fibril samples were deposited onto glow-discharged Quantifoil R 1.2/1.3 300-mesh copper grids, and excess sample was removed by blotting with filter paper prior to vitrification by plunge-freezing in liquid ethane using a Vitrobot Mark IV (FEI). Initial cryo-EM screening was conducted on either the Talos Arctica or Glacios instruments at the Cryo-Electron Microscopy Facility (CEMF), University of Texas Southwestern Medical Center (UTSW). Final high-resolution datasets were acquired on a 300 kV Titan Krios microscope (FEI) at two facilities: the CEMF and the Stanford-SLAC Cryo-EM Center (S2C2).

Raw movie frames underwent gain correction, alignment, motion correction, and dose weighting within RELION's integrated motion correction module[6]. Contrast transfer function (CTF) parameters were estimated using CTFFIND 4.1[7]. Helical reconstruction, three-dimensional (3D) refinement, and all post-processing steps were executed within the RELION 4.0 framework[8].

The filaments were picked automatically using Topaz in RELION 4.0. Initial particle extraction employed a box size of 1,024 pixels with an inter-box spacing of 15 asymmetric units (4.75 Å), and an additional extraction at 256 pixels with an inter-box spacing of 2 asymmetric units (4.75 Å). Helical parameters were estimated from 2D classification of the 1,024-pixel particle stack, while 2D classifications of the 256-pixel stack informed particle selection for downstream processing. The fibril helix was assumed to adopt a left-handed topology. To resolve distinct filament populations, 3D classifications were run with approximately 30,000–40,000 particles per class, using an elongated Gaussian blob as the starting reference model. Particle subsets yielding the highest-quality reconstructions were carried forward into 3D auto-refinement. Subsequent CTF refinement and Bayesian polishing steps were applied to push resolution further. Final maps were post-processed following standard RELION-recommended procedures. The curated particle subset was then used for high-resolution gold-standard refinement as previously described[10]. Overall resolution was assessed using the Fourier Shell Correlation (FSC) = 0.143 criterion between two independently refined half-maps.

**Model building**

The refined density map was sharpened using phenix.auto_sharpen. Previously deposited structural models of ATTRv-I84S (PDB: 8TDO and 8TDN) served as the template for constructing all near-atomic resolution models. Sequence mutations were introduced and iterative rigid body fitting and real-space refinement were carried out within COOT to arrive at the final models[11].

**Tryptic mass spectrometry (MS) sample preparation, data acquisition, and analysis**

0.5 µg of extracted ATTR fibrils were denatured by boiling in tricine SDS buffer and separated on a Novex™ 16% tris-tricine gel. Following Coomassie staining, the ATTR-associated smear was excised and subjected to overnight in-gel trypsin digestion. Resulting peptides were desalted by solid-phase extraction and analyzed by liquid chromatography-tandem mass spectrometry on a Q Exactive HF mass spectrometer interfaced with an Ultimate 3000 RSLC-Nano system. Precursor ions were scanned at 120,000 resolution, with up to 20 precursors per cycle selected for fragmentation by higher-energy collisional dissociation (HCD). Data were searched in Proteome Discoverer v3.0 SP1 (Thermo) using a semi-tryptic Sequest HT algorithm against the reviewed human UniProt database, allowing up to three missed cleavages, with precursor and fragment mass tolerances of 10 ppm and 0.02 Da, respectively. Cysteine carbamidomethylation was set as a fixed modification and methionine oxidation as a variable modification, with peptide identifications filtered at a 1% false discovery rate (FDR). All mass spectrometry proteomics data have been deposited to MassIVE with the accession number MSV000101672 [ftp://MSV000101672@massive-ftp.ucsd.edu].

### Reporting summary

Additional details on research design is available in the Nature Portfolio Reporting Summary associated with this article.

### Supplementary References


1. Nguyen, B. A. *et al.* Structural polymorphism of amyloid fibrils in ATTR amyloidosis revealed by cryo-electron microscopy. *Nat. Commun.* **15**, 581 (2024).
2. Fernandez-Ramirez, M. D. C. *et al.* Structural and molecular homogeneity of ATTRv-T60A amyloid fibrils across patients and organs. *Structure* **33**, 2013-2019.e4 (2025).
3. Nguyen, B. A. *et al.* Cryo-EM confirms a common fibril fold in the heart of four patients with ATTRwt amyloidosis. *Commun. Biol.* **7**, 905 (2024).
4. Nguyen, B. A. *et al.* ATTRv-V30M amyloid fibrils from heart and nerves exhibit structural homogeneity. *Structure* **32**, 2244-2250.e3 (2024).
5. Fernandez-Ramirez, M. D. C. *et al.* A shared amyloid fold in cardiac fibrils from three neuropathy-associated ATTR variants. *Structure* **34**, 487-495.e3 (2026).
6. Zivanov, J., Nakane, T. & Scheres, S. H. W. A Bayesian approach to beam-induced motion correction in cryo-EM single-particle analysis. *IUCrJ* **6**, 5–17 (2019).
7. Rohou, A. & Grigorieff, N. CTFFIND4: Fast and accurate defocus estimation from electron micrographs. *J. Struct. Biol.* **192**, 216–221 (2015).
8. He, S. & Scheres, S. H. W. Helical reconstruction in RELION. *J. Struct. Biol.* **198**, 163–176 (2017).
9. Scheres, S. H. W. RELION: Implementation of a Bayesian approach to cryo-EM structure determination. *J. Struct. Biol.* **180**, 519–530 (2012).
10. Emsley, P., Lohkamp, B., Scott, W. G. & Cowtan, K. Features and development of *Coot*. *Acta Crystallogr. D Biol. Crystallogr.* **66**, 486–501 (2010).
